# Detection of the HIV-1 accessory proteins Nef and Vpu by flow cytometry represents a new tool to study their functional interplay within a single infected CD4+ T cell

**DOI:** 10.1101/2021.11.03.467116

**Authors:** Jérémie Prévost, Jonathan Richard, Romain Gasser, Halima Medjahed, Frank Kirchhoff, Beatrice H. Hahn, John C. Kappes, Christina Ochsenbauer, Ralf Duerr, Andrés Finzi

**Author notes:** Contributed equally (Jérémie Prévost, Jonathan Richard). **Corresponding author: Andrés Finzi**, Centre de recherche du CHUM (CRCHUM), 900 St-Denis street, Tour Viger, R09.420, Montréal, Québec, H2X 0A9, Canada.

## Abstract

The HIV-1 Nef and Vpu accessory proteins are known to protect infected cells from antibody-dependent cellular cytotoxicity (ADCC) responses by limiting exposure of CD4-induced (CD4i) envelope (Env) epitopes at the cell surface. Although both proteins target the host receptor CD4 for degradation, the extent of their functional redundancy is unknown. Here, we developed an intracellular staining technique that permits the intracellular detection of both Nef and Vpu in primary CD4+ T cells by flow cytometry. Using this method, we show that the combined expression of Nef and Vpu predicts the susceptibility of HIV-1-infected primary CD4+ T cells to ADCC by HIV+ plasma. We also show that Vpu cannot compensate for the absence of Nef, thus providing an explanation for why some infectious molecular clones that carry a LucR reporter gene upstream of Nef render infected cells more susceptible to ADCC responses. Our method thus represents a new tool to dissect the biological activity of Nef and Vpu in the context of other host and viral proteins within single infected CD4+ T cells.

**IMPORTANCE:** HIV-1 Nef and Vpu exert several biological functions that are important for viral immune evasion, release and replication. Here, we developed a new method allowing simultaneous detection of these accessory proteins in their native form together with some of their cellular substrates. This allowed us to show that Vpu cannot compensate the lack of a functional Nef, which has implication for studies that use Nef-defective viruses to study ADCC responses.

## INTRODUCTION

The human immunodeficiency virus type 1 (HIV-1) genome encodes four accessory proteins (Vif, Vpr, Vpu and Nef), which are dispensable for viral replication *in vitro* but required for efficient replication, restriction factors counteraction and immune evasion *in vivo* (1–7). Among them, Nef and Vpu are well known for their role in subverting the host cell protein trafficking machinery (8, 9).

HIV-1 Nef is a small cytoplasmic protein of 27 kDa produced from early viral transcripts (10), which requires a myristoyl group on its N-terminus to traffic to intracellular and plasma membranes (11). Nef harbors a highly conserved dileucine motif in its C-terminal flexible loop that is responsible for the interaction with clathrin adaptors protein complexes (AP-1, AP-2 and AP-3) (12). Among these, interaction with AP-2 is required to downregulate the CD4 receptor from the surface of infected cells (13, 14) and target it for degradation in lysosomal compartments (15, 16).

HIV-1 Vpu is a small type-I transmembrane protein of 16 kDa produced late in the viral replication cycle (17, 18) and contains a short luminal N-terminal peptide followed by a single helical transmembrane domain and a C-terminal cytoplasmic domain (19–21). The cytoplasmic domain is comprised of two α-helices linked by a flexible loop known for its interaction with the SCF^βTRCP^ E3 ubiquitin ligase complex via a conserved phosphoserine motif (DS^P^GNES^P^) (22, 23). Vpu mainly localizes within intracellular compartments, notably the endoplasmic reticulum (ER) and the *trans*-Golgi network (TGN) (24–26). Like Nef, Vpu also induces degradation of newly synthesized CD4 by directing it through an ER-associated pathway (ERAD) for further proteasomal degradation (22, 27–29). In addition, Vpu sequesters the restriction factor BST-2 in the TGN using its transmembrane domain, thereby increasing the release of progeny virions (30–33).

CD4 downregulation by Nef and Vpu was previously reported to be critical for efficient viral replication in T cells by enhancing virion release and infectivity, and by preventing superinfection (34–39). CD4 downregulation is critical for immune evasion since the anti-Env antibody (Ab) response is dominated by non-neutralizing antibodies (nnAbs) that target Env in its “open” CD4-bound conformation (40–42). The interaction between CD4 and Env at the surface of HIV-1-infected cells has been shown to promote nnAbs binding to Env, leading to the elimination of infected cells through Fc-mediated effector functions, including antibody-dependant cellular cytotoxicity (ADCC) (41, 43). Nef and Vpu limit the presence of Env-CD4 complexes at the cell surface and thus protect infected cells against ADCC (41, 43, 44).

In previous studies, Nef and Vpu expression was mostly examined in transfected cell lines, frequently using tagged proteins (30, 31, 45, 46) or by performing Western blots and immunofluorescence microscopy in infected primary cells (47–52). However, both proteins are small, intracellularly located and present in low amounts, rendering their detection difficult. To facilitate their analysis in primary CD4+ T cells, we developed an intracellular staining technique to detect Nef and Vpu expression by flow cytometry, which allows the simultaneous detection of these proteins together with host and viral proteins within a single infected cell. Using this method, we show that Nef and Vpu expression predicts the susceptibility of HIV-1-infected primary CD4+ T cells to ADCC by HIV+ plasma. We also explain why decreased Nef expression in widely-used reporter viruses increase the susceptibility of infected cells to ADCC responses.

## RESULTS

### Intracellular detection of Nef and Vpu in HIV-1-infected primary CD4+ T cells

To facilitate detection of intracellular Nef, we obtained a polyclonal Nef antiserum through the NIH AIDS Reagent Program, which was generated by immunization of rabbits with a recombinant clade B Nef consensus protein produced in *E. coli* (53). In previous studies, this antibody detected native Nef proteins by Western blot and immunofluorescence microscopy in both transfected and infected cells (47, 54–56). Given the scarcity of anti-Vpu antibodies, we immunized rabbits with a peptide corresponding to the clade B Vpu C-terminal region (residues 69–81). A similar approach was previously used to generate a polyclonal antibody capable of detecting Vpu by Western blot and immunofluorescence microscopy (24, 57).

We first evaluated the ability of both Nef and Vpu antisera to recognize their cognate antigen using HEK 293T cells transfected with plasmids expressing the Nef or Vpu proteins from the transmitted/founder (T/F) virus CH058 (58, 59). Forty-eight hours post-transfection, cells were permeabilized and stained with the antisera, followed by detection with a fluorescently-labelled anti-rabbit secondary antibody. As expected, the Nef antiserum recognized only Nef transfected cells, while the Vpu antiserum recognized only Vpu transfected cells (Fig. 1A-C). To evaluate whether our method detected Nef and Vpu when expressed in a biologically relevant culture system, we infected primary CD4+ T cells with CH058 infectious molecular clones (IMC) encoding Nef, and/or Vpu proteins. While wildtype (WT)-infected cells were efficiently recognized by both Nef and Vpu antisera, abrogation of Nef (Fig 1D-E) or Vpu (Fig. 1F-G) expression prevented the recognition of productively-infected cells as identified by Gag protein intracellular staining (p24+). Of note, mock-infected or uninfected bystander cells (p24-) where not detected by either antiserum, further confirming their specificity (Fig 1D-G).

**Figure 1.**
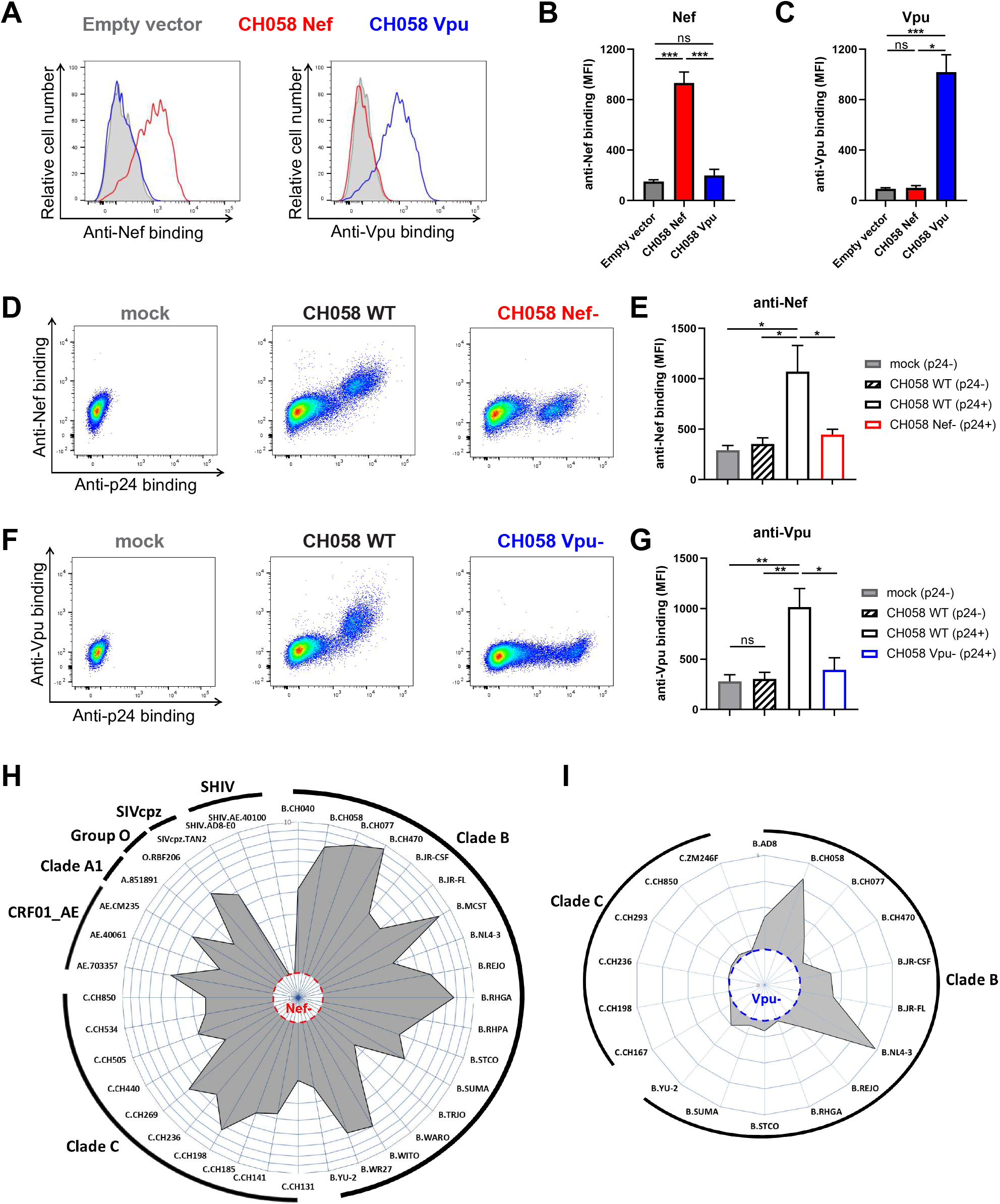
Intracellular detection of Nef and Vpu in infected primary CD4+ T cells. (**A-C**) 293T cells transfected with an empty vector or a plasmid expressing either CH058 Nef or CH058 Vpu. 48 hours post-transfection, cells were permeabilized and stained with rabbit polyclonal antisera raised against Nef and Vpu to detect their respective intracellular expression. Antiserum binding was detected using donkey anti-rabbit BV421 secondary Abs. (**A**) Histograms depicting representative staining and (**B-C**) Median fluorescence intensities (MFI) obtained for multiple independent stainings using (**B**) anti-Nef or (**C**) anti-Vpu. (**D-G**) Primary CD4+T cells mock-infected or infected with CH058 T/F WT, Nef- or Vpu-, were stained to detect the intracellular expression of Nef or Vpu. (**D,F**) Dot plots (left) and histograms (right) depicting representative (**D**) Nef and (**F**) Vpu staining. (**E,G**) The graphs show the MFI obtained from different cell populations using cells from five different donors using (**E**) anti-Nef or (**G**) anti-Vpu. Error bars indicate means ± standard errors of the means (SEM). Statistical significance was tested using an unpaired t test or a Mann-Whitney U test based on statistical normality (*, P < 0.05; **, P < 0.01; ***, P < 0.001; ns, nonsignificant). (**H-I**) Primary CD4+ T cells were infected with a panel of viruses from different clades (A1, B, C, CRF01_AE), group (M, O) and host (HIV-1, SIVcpz, SHIV). The radar plots indicate the level of specific recognition of infected cells (MFI normalized to uninfected cells) using the (**H**) anti-Nef or (**I**) anti-Vpu antisera. The limit of detection was determined using (**H**) cells infected with CH058 Nef- for Nef staining and using (**I**) cells infected with CH058 Vpu- for Vpu staining.

We next examined the antiserum binding to Nef and Vpu proteins from different HIV-1 clades and groups as well as from closely related simian immunodeficiency viruses (SIV). Primary CD4+ T cells were infected with a panel of HIV-1 IMCs representing clades B, C, A1 and CRF01_AE. As expected, both Nef and Vpu antisera recognized their respective antigen in cells infected with clade B viruses since both were raised against clade B immunogens (Fig. 1H-I). The anti-Nef polyclonal antibody was also able to recognize Nef proteins from group M clades C, A1 and CRF01_AE as well as the Nef from a group O isolate. This recognition extended even to the Nef protein of a related SIVcpzPts strain (isolate TAN2) but not to chimeric simian-human immunodeficiency viruses (SHIV) which express a SIVmac Nef (Fig. 1H). The Vpu antiserum was less cross-reactive and failed to detect Vpu from clade C viruses (Fig. 1I). These findings confirmed the specificity and cross-reactivity of the intracellular detection of Nef and Vpu using infected primary CD4+ T cells.

### Measuring CD4 and BST-2 downregulation in infected primary CD4+ T cells with or without Nef and Vpu expression

The efficient detection of Nef and Vpu at the single cell level by flow cytometry allowed us to combine this approach with the quantification of CD4 and BST-2 expression levels on the cell surface. Productively-infected cells (p24+) expressing both Nef and Vpu had little detectable CD4 and BST-2 compared to uninfected cells (Fig. 2A). In contrast, cells infected with Vpu or Nef mutant viruses differed in the extent of CD4 and BST-2 downregulation (Fig. 2A).

**Figure 2.**
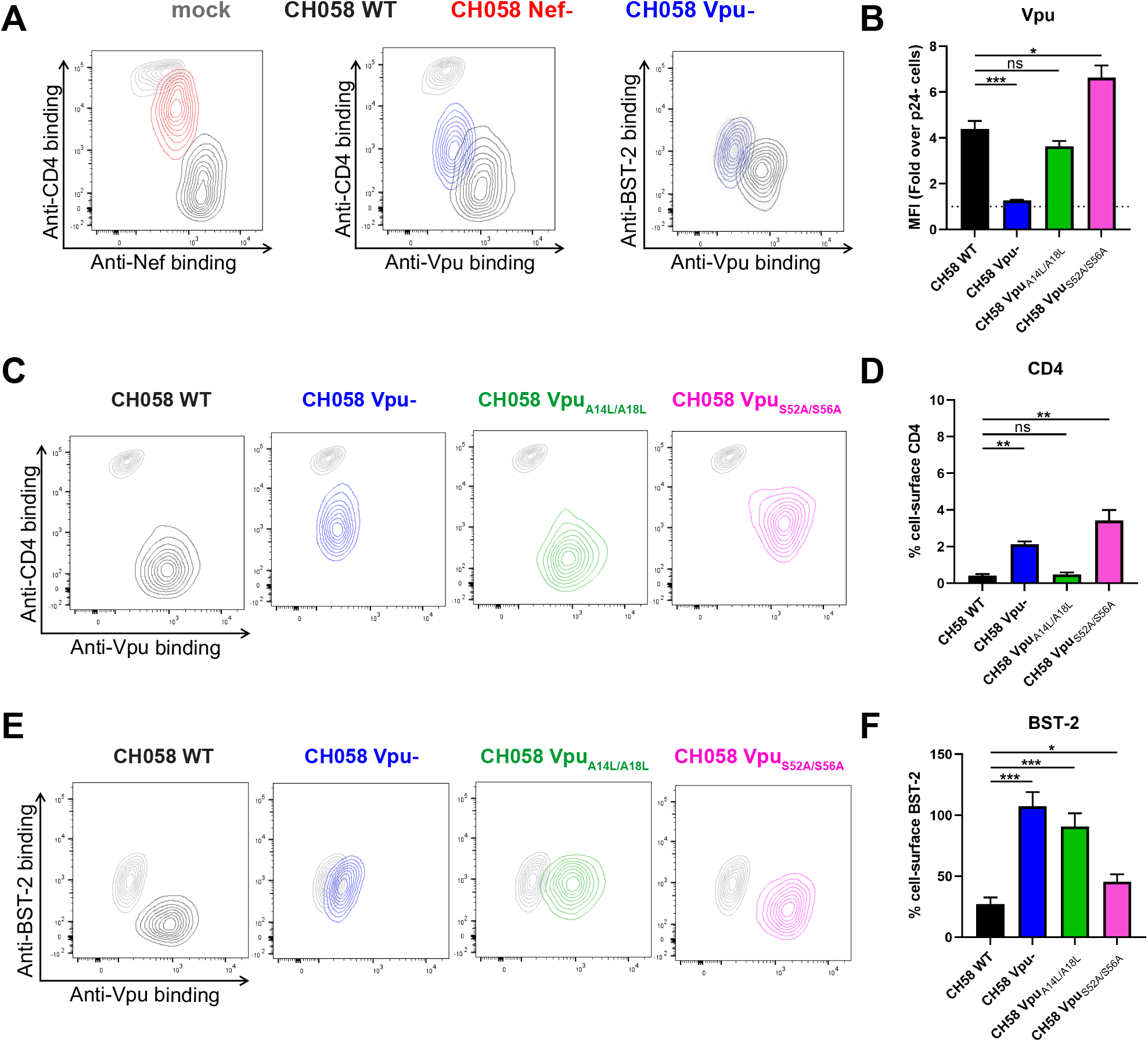
Concomitant detection of intracellular Nef and Vpu and cell-surface CD4 and BST-2. Primary CD4+T cells infected with CH058 T/F WT, Nef-, Vpu-, Vpu A14L/A18L or Vpu S52A/S56A viruses were stained for cell-surface CD4 and BST-2 prior to detection of intracellular Nef or Vpu expression. (**A,C,E**) Contour plots depicting representative cell-surface CD4 or BST-2 detection in combination with Nef or Vpu intracellular detection. Mock-infected cells were used as a control and are shown in grey. (**B,D,F**) The graphs show MFIs obtained from five independent experiments. Error bars indicate means ± standard errors of the means (SEM). Statistical significance was tested using an unpaired t test or a Mann-Whitney U test based on statistical normality (*, P < 0.05; **, P < 0.01; ***, P < 0.001; ns, nonsignificant).

Vpu targets CD4 and BST-2 by different mechanisms. First, Vpu interacts with multiple transmembrane proteins, including BST-2, through its transmembrane domain (TMD), which sequesters these proteins in perinuclear compartments (32, 33, 60–63). Second, Vpu downregulates CD4 by interaction of its cytoplasmic domain with the cytoplasmic tail of CD4 (64–69). Consistent with these different interaction modes, Vpu-mediated CD4 and BST-2 degradation involves independent pathways (proteasomal and lysosomal degradation, respectively), both of which depend on polyubiquitination by the SCF^βTRCP^ E3 ubiquitin ligase complex, recruited by Vpu using its highly conserved phosphoserine motif (22, 26, 70, 71). To examine whether we could measure the expression and activity of Vpu mutants by flow cytometry, we introduced mutations at critical residues of the Vpu TMD (A14L/A18L) or its phosphoserine motif (S52A/S56A). CH058 IMCs coding for wildtype or mutated Vpu proteins were used to infect primary CD4+ T cells. While the TMD mutations did not affect Vpu expression, the phosphoserine mutations led to a significant accumulation of intracellular Vpu proteins (Fig. 2B), most likely because Vpu is degraded together with its target protein as a ubiquitinated complex (24, 72, 73). Despite a higher expression, the Vpu phosphoserine mutant was unable to downregulate CD4 and marginally diminished in its capacity to antagonize BST-2 (Fig 2C-F). This is consistent with studies demonstrating that the recruitment of the SCF^βTRCP^ E3 ubiquitin ligase complex and the degradation of BST-2 by Vpu is dissociable from its capacity to antagonize the restriction factor (32, 71, 74–76). In contrast, the Vpu TMD mutations did not affect Vpu’s ability to target CD4 but completely abrogated its capacity to downregulate BST-2 (Fig 2C-F). Together, these results emphasize the need of measuring Nef and Vpu expression when studying their biological functions.

### Nef and Vpu expression inversely correlates with ADCC responses

CD4 downregulation by Nef and Vpu, together with Vpu-mediated BST-2 antagonism were found to be critical factors preventing the exposure of vulnerable CD4-induced Env epitopes, thus protecting HIV-1-infected cells from ADCC (41, 43, 44, 77–80). To investigate the link between Nef and Vpu expression and HIV-1-infected cells immune evasion, we infected activated primary CD4+ T cells from five HIV-negative individuals with two clade B IMCs, CH058 T/F and JR-FL, encoding functional or defective *nef* and *vpu* genes. Focusing on the productively-infected cells (p24+), we performed a comprehensive characterization of the patterns of viral protein expression including cell-surface Env (detected with the conformation-independent Ab 2G12), intracellular Nef and Vpu in combination with cell-surface levels of CD4 and BST-2. We also measured the specific recognition and elimination of infected cells by ADCC using the CD4-induced (CD4i) A32 monoclonal Ab (mAb). This antibody binds the cluster A region of the gp120 which is occluded in the “closed” trimer and therefore can only bind Env in the “open” CD4-bound conformation. We also tested 25 different plasma samples from chronically HIV-1-infected individuals. As expected, Nef was only expressed by WT and Vpu-constructs, while Vpu was only expressed by WT and Nef-constructs (Fig. 3A). Consistent with previous reports (43, 77, 78), deletion of Nef strongly impaired CD4 downregulation by both viruses but did not affect Env or BST-2 cell-surface levels. Vpu deletion mitigated CD4 downregulation to a lesser extent than Nef and abrogated BST-2 downmodulation, resulting in an overall increase in the amount of cell-surface Env (Fig. 3B). We noticed lower levels of the JR-FL Vpu protein compared to CH058 Vpu, which was linked to a less effective Vpu-mediated CD4 downmodulation (Fig. 3B). The cumulative effect of Nef and Vpu on cell-surface levels of Env, CD4 and BST-2 prevented the recognition of infected cells and protected them from ADCC responses mediated by A32 and HIV+ plasma (Fig. 3C-D). In contrast, abrogation of Nef or Vpu expression resulted in increased recognition and susceptibility of infected cells to ADCC mediated by nnAbs (Fig. 3C-D). We performed correlation analyses to measure the level of association between the different cellular, virological and immunological variables (Fig 3E-F). We found that both Nef and Vpu established a large network of inverse correlations with cellular and immunological factors. Interestingly, Env levels hardly contributed to the network and were poorly associated with the immunological outcome, thus indicating that the overall amount of Env present at the surface does not dictate ADCC responses mediated by CD4i Abs or HIV+ plasma, but rather the conformation Env occupies. Apart from antibody binding, ADCC responses mediated by nnAbs correlated strongly with CD4 and Nef levels (Fig. 3E-F). Overall, Nef and Vpu expression inversely correlates with the susceptibility of HIV-1-infected cells to ADCC mediated by CD4i Abs and HIV+ plasma.

**Figure 3.**
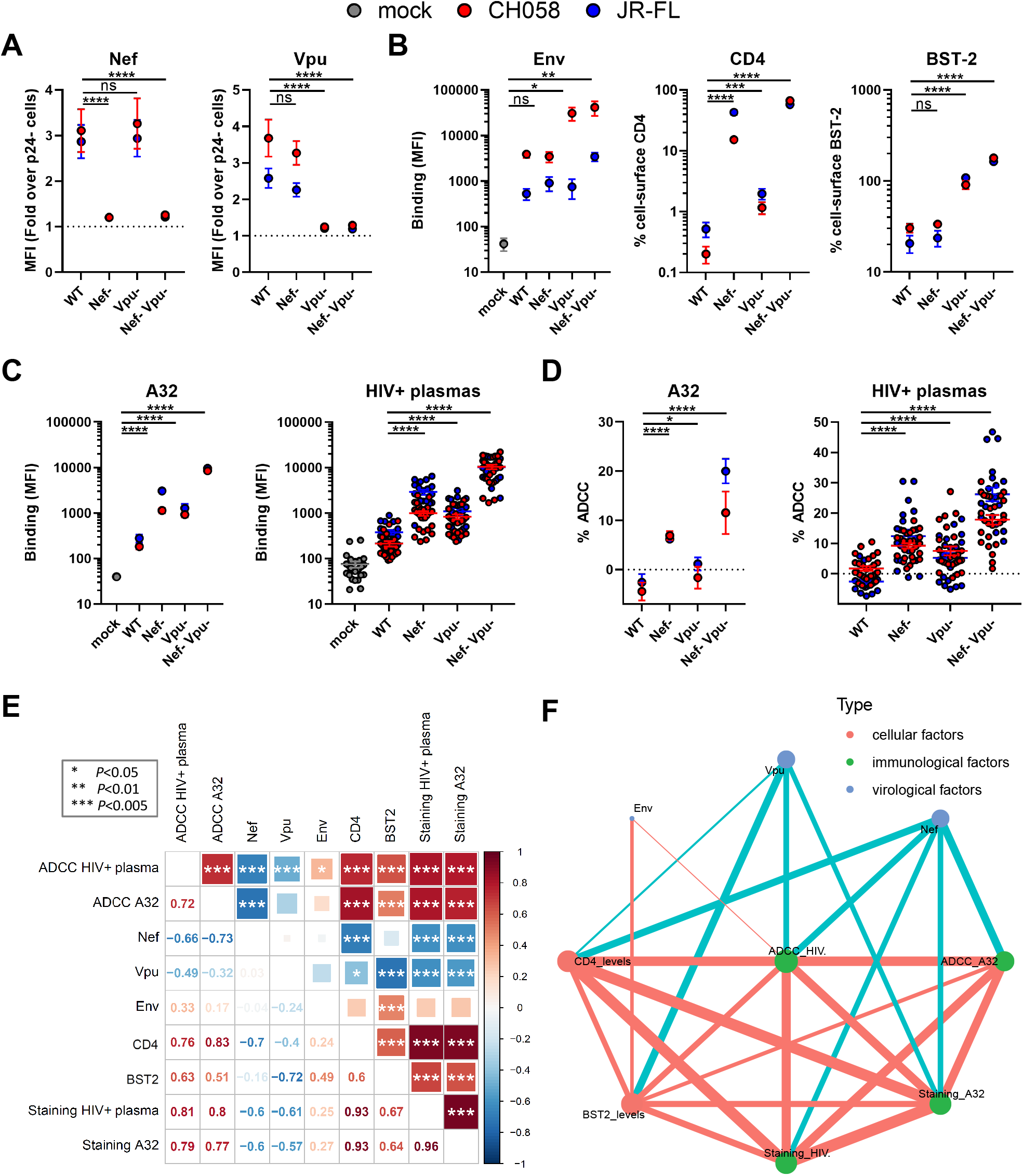
Nef and Vpu intracellular detection inversely correlates with the recognition of infected cells and their susceptibility to ADCC responses mediated by HIV+ plasma. Primary CD4+T cells were mock-infected (grey), infected with CH058 T/F (red) or JR-FL (Blue) viruses (WT, Nef-, Vpu-, Nef-Vpu-) and stained for (**A**) intracellular Nef or Vpu expression in combination with (**B**) cell-surface staining of Env (using the anti-Env 2G12 mAb), CD4 and BST-2. The ability of the anti-Env A32 mAb and 25 different HIV+ plasma to (**C**) recognize infected cells and (**D**) eliminate infected cells by ADCC was also measured. (**A-D**) The graphs show the MFI obtained on the infected (p24+) cell population using cells from five different donors. Error bars indicate means ± standard errors of the means (SEM). Statistical significance was tested using an unpaired t test or a Mann-Whitney U test based on statistical normality (*, P < 0.05; **, P < 0.01; ***, P < 0.001; ns, nonsignificant). (**E**) Correlograms summarize pairwise correlations among all immunological, virological and cellular variables obtained from infected primary CD4+ T cells (shown in A-D). Squares are color-coded according to the magnitude of the correlation coefficient (r) and the square dimensions are inversely proportional with the P-values. Red squares represent a positive correlation between two variables and blue squares represent negative correlations. Asterisks indicate all statistically significant correlations (*P < 0.05, **P < 0.01, ***P < 0.005). Correlation analysis was done using nonparametric Spearman rank tests. (**F**) Correlation networks were generated using data shown in (**E**). Each node (circle) represents a cellular (red), an immunological (green) or a virological (blue) feature measured on infected cells. Nodes are connected with edges (lines) if they are significantly correlated (P < 0.05); nodes without edges were removed. Edges are weighted according to P-values (inversely). Red edges represent a positive correlation between two variables and blue edges represent negative correlations. Nodes are sized according to the r-values of connecting edges.

### Impaired Nef expression from IMC LucR.T2A constructs enhance the susceptibility of infected cells to ADCC

Infectious molecular clones encoding for the *Renilla* luciferase (LucR) reporter gene upstream of the *nef* sequence, and a T2A ribosome-skipping peptide to drive Nef expression are widely employed to quantify anti-HIV-1 ADCC responses (81–92). Despite evidence that Nef-mediated CD4 downregulation is impaired when using these IMCs (54, 79), a series of recent studies have hypothesized that Vpu can compensate for the absence of Nef expression and completely downregulate CD4 on its own (93–97). To evaluate this hypothesis, we used our intracellular staining to measure Nef and Vpu expression levels and study their impact on ADCC responses mediated by nnAbs against cells infected with IMC-LucR.T2A constructs. Primary CD4+ T cells were infected with NL4.3-based IMCs that do (Env-IMC-LucR.T2A) or do not encode (Env-IMC) a LucR.T2A cassette. These IMCs express the Env ectodomain from two clade B viruses, CH058 T/F and YU-2. Consistent with the lack of Nef detection by Western blot (54, 79), insertion of the LucR.T2A cassette also impaired the detection of Nef by flow cytometry, while Vpu expression remained unchanged (Fig. 4A-B). However, we noted an accumulation of cell-surface CD4 for Env-IMC-LucR.T2A compared to *nef*-intact constructs (~20-fold higher) (Fig. 4C), which resulted in a significantly increased recognition and susceptibility of infected cells to ADCC responses mediated by A32 mAb and HIV+ plasma (Fig. 4D-E). Of note, both the binding and the ADCC responses mediated by nnAbs were strongly associated with CD4 levels and inversely correlated with Nef expression (Fig 4F-G). In contrast, these variables poorly correlated with Vpu expression. Based on these data, it seems clear that Vpu expression alone is not sufficient to prevent ADCC-mediated killing of infected cells and that HIV-1 requires both Nef and Vpu for efficient humoral response evasion.

**Figure 4.**
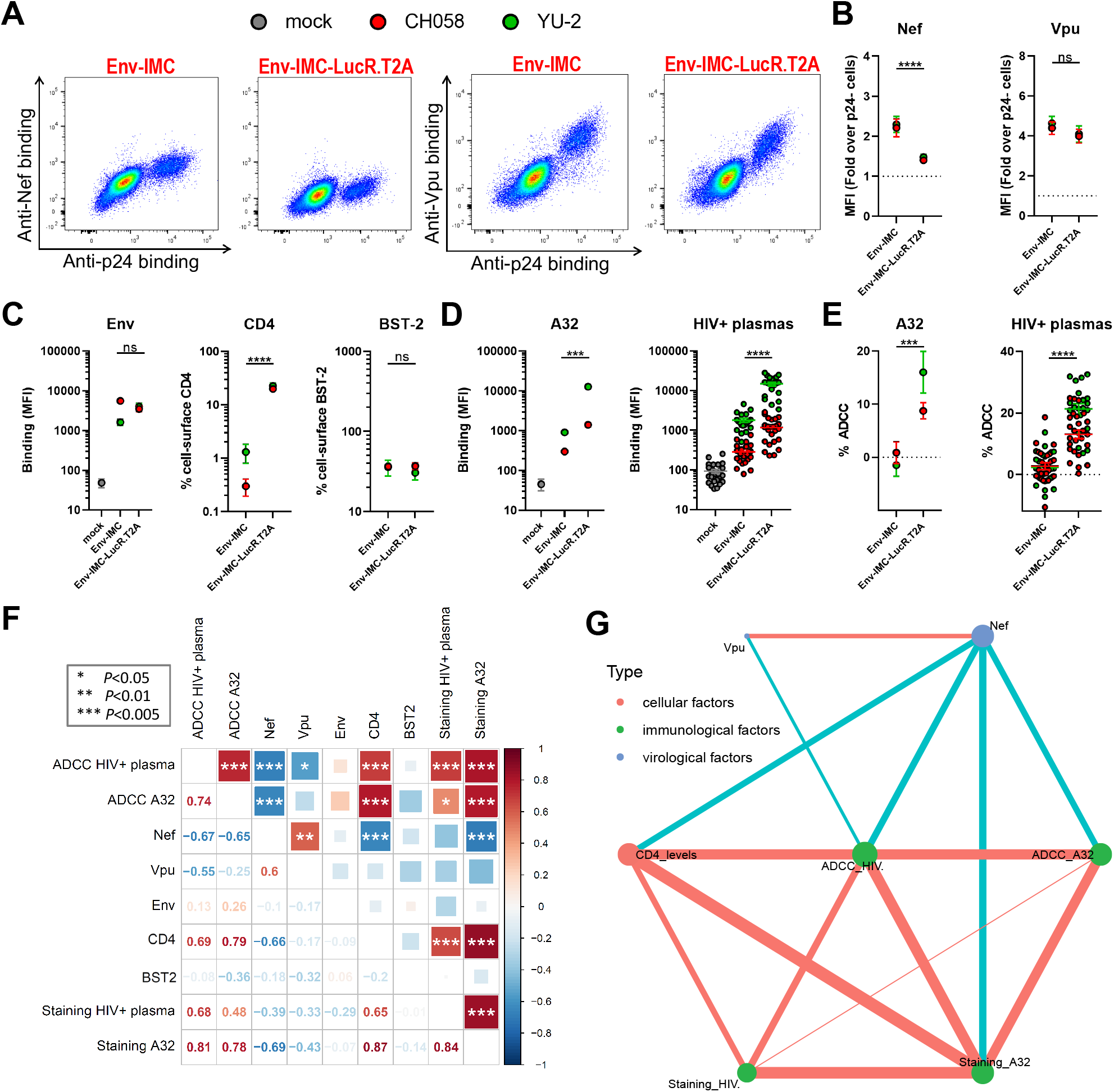
Lack of Nef expression in primary CD4+ T cells infected with LucR.T2A IMCs results in enhanced ADCC. Primary CD4+T cells mock-infected (grey) or infected with chimeric IMCs expressing CH058 Env (red) or YU-2 Env (green) and expressing or not the LucR reporter gene. (**A**) Dot plots depicting representative stainings of intracellular Nef or Vpu expression. (**B-C**) Detection by flow cytometry of (**B**) intracellular Nef or Vpu expression in combination with (**C**) cell-surface staining of Env (using anti-Env mAbs 2G12 (CH058) or PGT135 (YU-2)), CD4 and BST-2. The ability of the A32 mAb and 25 HIV+ plasma to (**D**) recognize infected cells and (**E**) eliminate infected cells by ADCC was also measured. (**B-E**) The graphs show the MFI obtained on the infected (p24+) cell population using cells from five different donors. Error bars indicate means ± standard errors of the means (SEM). Statistical significance was tested using an unpaired t test or a Mann-Whitney U test based on statistical normality (*, P < 0.05; **, P < 0.01; ***, P < 0.001; ns, nonsignificant). (**F**) Correlograms summarize pairwise correlations among all immunological, virological and cellular variables obtained from infected primary CD4+ T cells (shown in **B-E**). Squares are color-coded according to the magnitude of the correlation coefficient (r) and the square dimensions are inversely proportional with the P-values. Red squares represent a positive correlation between two variables and blue squares represent negative correlations. Asterisks indicate all statistically significant correlations (*P < 0.05, **P < 0.01, ***P < 0.005). Correlation analysis was done using nonparametric Spearman rank tests. (**G**) Correlation networks were generated using data shown in (**F**). Each node (circle) represents a cellular (red), an immunological (green) or a virological (blue) feature measured on infected cells. Nodes are connected with edges (lines) if they are significantly correlated (P < 0.05); nodes without edges were removed. Edges are weighted according to P-values (inversely). Red edges represent a positive correlation between two variables and blue edges represent negative correlations. Nodes are sized according to the r-values of connecting edges.

### Nef, Vpu and CD4 levels predict ADCC responses mediated by HIV+ plasma

We next used univariate multiple linear regression (MLR) analysis to evaluate the capacity of different variables to predict ADCC responses mediated by HIV+ plasma. This model is based on the hypothesis that a linear relationship exists between the dependant variable quantified empirically and the independent variables that serve as predictive variables. In our model, the dependant variable is the ADCC responses mediated by plasma from HIV+ donors (ADCC HIV+ plasma) and the independent variables are the cellular, virological and immunological factors measured on infected cells. To run the MLR model, we combined data obtained with the different viral constructs (Fig. 3 & 4) and plotted the mean ADCC obtained with 25 HIV+ plasma against a single virus on the X axis and the associated predicted ADCC value based on one or more independent variables on the Y axis. When looking at cellular factors, we noticed that only CD4 accurately predicts ADCC responses mediated by HIV+ plasma, independent of the viral strain (Fig. 5A). Even though BST-2 displayed a strong correlation with ADCC responses (Fig 3E), it was not predictive. When focusing on virological variables, we observed that Nef is the only significant ADCC predictive variable, albeit not as good as CD4 (Fig. 5A-B). However, combinations of Nef with Vpu or Env increased its predictive scores, reaching similar levels as CD4 when combined with Vpu (Fig. 5B). Of note, the strength of the prediction was not further improved when combining all three virological variables altogether. As for immunological variables, their capacity to predict ADCC by HIV+ plasma was found to be equivalent or even higher than for cellular and virological factors (Fig. 5C). Indeed, the binding of HIV+ plasma predicted ADCC values with a similar score as CD4 or Nef and Vpu combined, while the binding of A32 predicted ADCC by HIV+ plasma even better (Fig. 5A-C). This could be explained by the fact that A32-like Abs present in plasma from infected individuals are from the main class of Abs (anti-cluster A Abs) mediating ADCC responses against infected cells (41, 80, 81, 91, 98). In line with this interpretation, ADCC mediated by A32 was found to have a near-perfect predictive ability, suggesting that factors, other than antibody binding, are presumably needed to fully explain the ADCC phenotypes observed (Fig. 5C).

**Figure 5.**
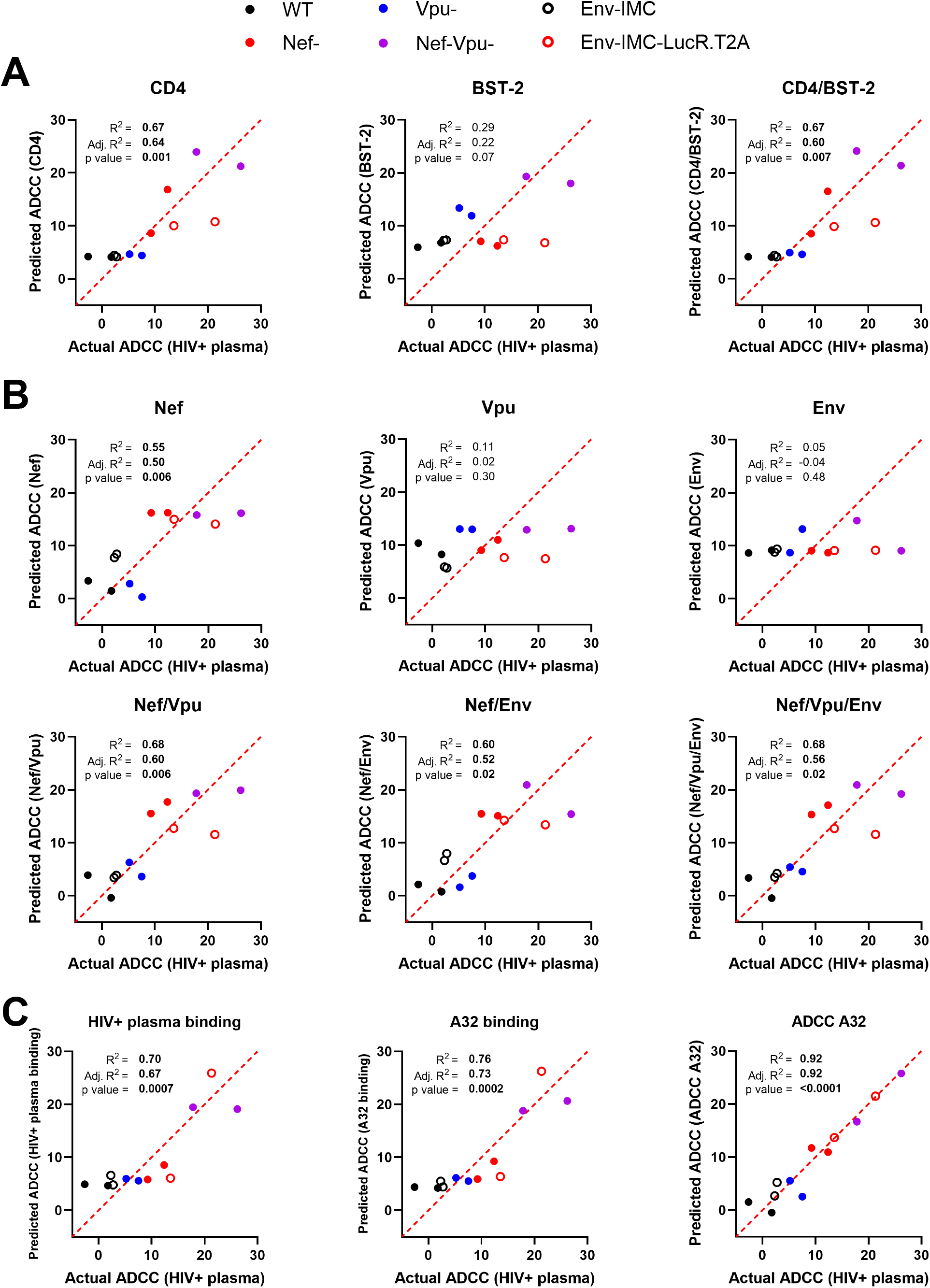
Prediction of ADCC responses mediated by HIV+ plasma using multiple linear regression models. (**A-C**) Multiple linear regression analysis to identify variables that can predict the ADCC responses mediated by HIV+ plasma against primary CD4+ T cells infected by different viral constructs (WT, Nef-, Vpu-, Nef-Vpu-, Env-IMC, Env-IMC-LucR.T2A) from different HIV-1 strains (CH058, JR-FL, YU-2). Each dot represents a single virus where the average of ADCC obtained with 25 different HIV+ plasma (dependent variable) is plotted on the X axis and the predicted ADCC value based on one or more independent parameters is plotted on the Y axis. Predictors include (**A**) cellular variables, (**B**) virological variables and (**C**) immunological variables. Multiple linear regression analyses were performed using the GraphPad Prism software (version 9.1.0). P values below 0.05 are considered significant and are highlighted in bold. The coefficient of multiple correlation (R^2^) indicates the goodness of fit of the multiple regression linear model. The adjusted R^2^ (Adj. R^2^) is used to compare the fits of models across experiments with different numbers of data points and independent variables.

## DISCUSSION

Unlike simple retroviruses, HIV-1 and related SIVs encode multiple accessory proteins that promote viral replication and immune evasion (99). Among them, Nef and Vpu modulate the expression, trafficking, localization and function of several host cell surface proteins, including the viral receptor CD4, restriction factors and homing receptors (28, 30, 31, 62, 69, 100–104). They also modulate a wide range of immunoreceptors to evade immune responses mediated by CD8+ T, NK and NKT cells (105–113). Most of these host cell proteins are naturally expressed on primary CD4+ T cells, the preferential target of HIV-1. The detection of Nef and Vpu has previously been done in transfected cells (30, 31, 46, 48, 49, 114), which results in the overexpression of the viral proteins when compared to infected primary CD4+ T cells. Moreover, tagged viral proteins are frequently used to facilitate their detection (30, 31, 46, 48, 49, 114). Protein overexpression and/or tag insertion, have the potential to impact the trafficking and functions of these accessory proteins. To study Nef and Vpu’s biological activities in a physiologically relevant system, we developed an intracellular staining method to detect native Nef and Vpu proteins in HIV-1-infected primary CD4+ T cells by flow cytometry. Using Nef and Vpu antisera, we detected both viral proteins with high specificity in cells productively infected (p24+) with multiple IMCs. The Nef antiserum was cross-reactive, detecting Nef from group M (clade B, C, A1 and CRF01_AE), from a group O isolate and from a closely related SIVcpz strain. In contrast, the Vpu antiserum recognized only clade B Vpu proteins, consistent with the fact that we used a peptide from the C-terminal region of clade B Vpu. This region is highly variable among group M viruses (115). More conserved regions of Vpu map to the transmembrane domain of the protein and the βTRCP binding site (116, 117). However, these regions are either buried into the plasma membrane or occluded by cellular partners, and thus are not readily accessible for antibody recognition. While the generation of a broad Vpu antiserum is challenging, it may be possible to generate clade-specific Vpu antisera by immunization using peptides corresponding to the C-terminal region specific for a given clade.

Nef and Vpu intracellular detection by flow cytometry represents an excellent tool to study their biological activities in HIV-1-infected primary CD4+ T cells. This method allows for the detection of cell-surface substrates or antibody recognition of surface Env and the concomitant detection of Nef and Vpu expression within a single infected cell (Fig. 2A). Infected CD4+ T cells represent the most relevant system to study the complex interplay between these two accessory proteins and the wide range of host cell factors naturally expressed by T cells. Recent findings revealed that modulation of BST-2 levels by type I IFN impacts the capacity of Vpu to downregulate NTB-A, PVR, CD62L and Tim-3, thus reducing its polyfunctionality (63, 69). Nef and Vpu also display overlapping functions, as they share the capacity to downregulate several cell-surface proteins, including CD4, PVR, CD62L and CD28 (8, 56, 62, 110, 118). The expression levels of one viral protein could therefore modulate the biological activities of the other, making it essential to study their functions in a context where both viral proteins are expressed simultaneously at physiological levels. Thus, our intracellular staining measuring Nef/Vpu expression and functionality in HIV-1-infected cells represents a new approach to better characterize their functional interplay.

Increasing evidence points towards Env conformation on the surface of infected cells as a critical parameter of ADCC susceptibility to HIV+ plasma (41, 119–121). Non-neutralizing antibodies in the plasma from HIV-1-infected individuals target epitopes that are only exposed when Env interacts with cell-surface CD4, thus adopting the “open” CD4-bound conformation (41, 43). Nef and Vpu contribute to protect HIV-1-infected cells from ADCC by limiting Env-CD4 interaction via CD4 downregulation and BST-2 antagonism (41, 43, 44, 77, 78). Here we confirm and extend previous observations by showing that Nef and Vpu expression predicts the susceptibility of HIV-1-infected primary CD4+ T cells to ADCC responses. In agreement with recent studies (44, 79), we found that CD4 accurately predicted the susceptibility of infected cells to ADCC (Fig. 5). Given its enhanced capacity to downregulate CD4 compared to Env or Vpu (34, 41, 43, 118), Nef represents the main viral factor influencing ADCC responses mediated by CD4-induced ligands (Fig. 5B). On the contrary, BST-2 and Env expression, alone or in combination, were unable to accurately predict the susceptibility of infected cells to ADCC. These results are consistent with previous reports suggesting that Env conformation rather than overall cell-surface Env levels, drives ADCC responses mediated by HIV+ plasma (41, 43, 120, 121). This is also in agreement with recent work showing that BST-2 upregulation by type I IFN enhances cell-surface Env levels without increasing the susceptibility of infected cells to ADCC mediated by HIV+ plasma, unless CD4-mimetics are used to “open-up” Env and stabilize the CD4-bound conformation (122).

A series of recent studies using LucR.T2A IMCs have hypothesized that Vpu can compensate for the absence of Nef expression by fully downregulating cell-surface CD4 (93–97). Our results show that this is not the case. Consistent with its role in targeting CD4 already present at the plasma membrane, the impact of Nef on CD4 downmodulation is more prominent (Fig. 2 & 3) (34, 41, 43, 118). In its absence, Vpu was unable to fully downregulate CD4, thus sensitizing infected cells to ADCC responses. These results highlight the importance of selecting full-length unmutated IMCs with proper Nef and Vpu expression to generate biologically relevant ADCC measurements. For example, a recent manuscript recently reported no differences in ADCC susceptibility between cells infected with clade B, clade C or CRF01_AE IMCs (123) while previous studies have shown otherwise (120, 124). In this article (123), the authors use functionally Nef defective LucR.T2A IMCs, which results in incomplete CD4 downregulation and therefore exposure of Env in its CD4-bound conformation at the cell surface (Fig. 4) (79). Thus, it is not surprising that the usage of Nef defective viruses skew ADCC responses in favor of nnAbs and mitigate the intrinsic differences that exists between Env from different clades. Fortunately, several alternatives to the use of LucR.T2A IMCs are available to measure ADCC against productively-infected cells (125), including the Infected Cell Elimination (ICE) assay, which measures the loss of productively-infected cells (p24+) by flow cytometry and allows the utilization of unmodified IMCs. Utilization of an NK cells resistant T cell line expressing a Tat-driven luciferase reporter gene (CEM.NKr-CCR5-sLTR-Luc) as target cells also represents an option (126). Finally, luciferase reporter IMCs (referred to as LucR.6ATRi IMCs) expressing similar levels of Nef than those obtained with unmodified IMC are also available. These IMCs utilize a modified encephalomyocarditis virus (EMCV) internal ribosome entry site (IRES) element in lieu of T2A (54, 79). Thus, LucR.6ATRi reporter viruses represent a biologically relevant alternative to LucR.T2A IMCs when measuring ADCC mediated by nnAbs and plasma collected from infected or vaccinated individuals.

## ACKNOWLEDGMENTS

The authors thank the CRCHUM BSL3 and Flow Cytometry Platforms for technical assistance, Mario Legault from the FRQS AIDS and Infectious Diseases network for cohort coordination and clinical samples. We thank the following collaborators for kindly providing some infectious molecular clones: Dennis Burton (The Scripps Research Institute) for JR-FL, Malcom A. Martin (NIAID) for SHIV_AD8-EO_, George M. Shaw (UPenn) for SHIV.AE.40100 and Sodsai Tovanabutra (US MHRP) for the HIV-1_WR27_, HIV-1703357, HIV-140061, HIV-1_CM235_ and HIV-1851891 IMCs. We thank MédiMabs for their scientific and technical support during the development of the Vpu antiserum. This study was supported by grants from the National Institutes of Health to A.F., C.O. and J.C.K. (R01 AI148379), to A.F. (R01 AI129769 and R01 AI150322) and to BHH (R01 AI162646 and UM1 AI164570). This work was also partially supported by 1UM1AI164562-01, co-funded by National Heart, Lung and Blood Institute, National Institute of Diabetes and Digestive and Kidney Diseases, National Institute of Neurological Disorders and Stroke, National Institute on Drug Abuse and the National Institute of Allergy and Infectious Diseases, a CIHR foundation grant #352417, a CIHR Team grant #422148 and a Canada Foundation for Innovation grant #41027 to A.F. A.F. is the recipient of a Canada Research Chair on Retroviral Entry #RCHS0235 950-232424. F.K. is supported by the Deutsche Forschungsgemeinschaft (CRC 1279 and SPP 1923). J.P. is the recipient of a CIHR doctoral fellowship. The funders had no role in study design, data collection and analysis, decision to publish, or preparation of the manuscript.

## AUTHOR CONTRIBUTIONS

J.P. J.R. and A.F. conceived the study. J.P., J.R., and A.F. designed experimental approaches. J.P., J.R., R.G. R.D. and A.F. performed, analyzed, and interpreted the experiments. H.M., F.K., B.H.H., J.C.K. and C.O. supplied novel/unique reagents. J.P., J.R., BHH, and A.F. wrote the paper. All authors have read, edited, and approved the final manuscript.

## CONFLICT OF INTEREST

The authors declare no competing interests.

## DATA AVAILABILITY

Data and reagents are available upon request.

## METHODS

### Ethics Statement

Written informed consent was obtained from all study participants [the Montreal Primary HIV Infection Cohort (127, 128) and the Canadian Cohort of HIV Infected Slow Progressors (129–131), and research adhered to the ethical guidelines of CRCHUM and was reviewed and approved by the CRCHUM institutional review board (ethics committee, approval number CE 16.164 - CA). Research adhered to the standards indicated by the Declaration of Helsinki. All participants were adult and provided informed written consent prior to enrolment in accordance with Institutional Review Board approval.

### Cell lines and isolation of primary cells

HEK293T human embryonic kidney cells (obtained from ATCC) were grown as previously described (132). Primary human PBMCs and CD4+ T cells were isolated, activated and cultured as previously described (43). Briefly, PBMCs were obtained by leukapheresis from HIV-negative individuals (4 males and 1 female) and CD4+ T lymphocytes were purified from resting PBMCs by negative selection using immunomagnetic beads per the manufacturer’s instructions (StemCell Technologies, Vancouver, BC) and were activated with phytohemagglutinin-L (10 μg/mL) for 48 hours and then maintained in RPMI 1640 complete medium supplemented with rIL-2 (100 U/mL).

### Plasmids and proviral constructs

The vesicular stomatitis virus G (VSV-G)-encoding plasmid was previously described (133). Transmitted/Founder (T/F) and chronic infectious molecular clones (IMCs) of patients CH040, CH058, CH077, CH131, CH141, CH167, CH185, CH198, CH236, CH269, CH293, CH440, CH470, CH505, CH534, CH850, CM235, MCST, REJO, RHGA, RHPA, STCO, SUMA, TRJO, WARO, WITO, WR27, 40061, 703357 and 851891 were inferred, constructed, and biologically characterized as previously described (120, 134–143). The IMCs encoding for HIV-1 reference strains AD8, JR-FL, JR-CSF, NL4-3, YU-2 were described elsewhere (144–149). HIV-1 group O (RBF206), SIVcpz (TAN2) and chimeric SIVmac/HIV-1 IMC constructs (SHIV_AD8-EO_ and SHIV.AE.40100) were generated as previously published (150–153). CH058 IMCs defective for Vpu and/or Nef expression were previously described (58). To generate a *nef*-defective JR-FL IMC, a frameshift mutation was introduced at the unique XhoI restriction site within the *nef* gene, resulting in a premature stop codon at position 47. To generate *vpu*-defective JR-FL IMCs, two stop-codons were introduced directly after the start-codon of *vpu* using the QuikChange II XL site-directed mutagenesis protocol (Agilent Technologies, Santa Clara, CA). The presence of the desired mutations was determined by automated DNA sequencing. Proviral constructs, collectively referred as Env-IMCs, comprising an HIV-1 NL4.3-based isogenic backbone engineered for the insertion of heterologous *env* strain sequences and expression in *cis* of full-length Env (pNL.CH058.ecto and pNL.YU-2.ecto), were previously described (47). In the same study, isogenic proviral constructs encoding *Renilla* luciferase (LucR) followed in frame by a ribosome-skipping T2A peptide intended to drive Nef expression were also reported (collectively referred to as Env-IMC-LucR.T2A) (47). Construction of plasmids encoding for CH058 Vpu and CH058 Nef in the pCGCG-IRES-eGFP expression vector was previously described (58, 59).

### Viral production and infections

To achieve a similar level of infection in primary CD4^+^ T cells among the different IMCs tested, VSV-G-pseudotyped HIV-1 viruses were produced and titrated as previously described (120). Viruses were then used to infect activated primary CD4+ T cells from healthy HIV-1 negative donors by spin infection at 800 × *g* for 1 h in 96-well plates at 25 °C.

### Antibodies and plasma

The following Abs were used to assess cell-surface Env staining: A32, 2G12 (NIH AIDS Reagent Program) and PGT135 (IAVI). Mouse anti-human CD4 (clone OKT4; Thermo Fisher Scientific, Waltham, MA, USA) and mouse anti-human BST-2 (clone RS38E, PE-Cy7-conjugated; Biolegend, San Diego, CA, USA) were also used as primary antibodies for cell-surface staining. Goat anti-mouse and anti-human antibodies pre-coupled to Alexa Fluor 647 (Invitrogen, Rockford, IL, USA) were used as secondary antibodies in flow cytometry experiments. Plasma from HIV-infected individuals were collected, heat-inactivated and conserved at −80 °C until use. Rabbit antisera raised against a Nef consensus protein (NIH AIDS Reagent Program) or against a Vpu C-terminal peptide (69) were used as primary antibodies in intracellular staining. BrillantViolet 421 (BV421)-conjugated donkey anti-rabbit antibodies (Biolegend) were used as secondary antibodies to detect Nef and Vpu antisera binding by flow cytometry. To avoid any potential cross-reactivity with the anti-rabbit secondary antibodies used for intracellular staining, mouse monoclonal antibodies were used to detect CD4 and BST-2 proteins.

### Flow cytometry analysis of cell-surface and intracellular staining

Cell-surface staining of infected cells was performed as previously described (41). Binding of cell-surface HIV-1 Env by anti-Env mAbs (5 μg/mL) or HIV+ plasma (1:1000 dilution) was performed at 48h post-infection. Infected cells were then permeabilized using the Cytofix/Cytoperm Fixation/ Permeabilization Kit (BD Biosciences, Mississauga, ON, Canada) and stained intracellularly using PE-conjugated mouse anti-p24 mAb (clone KC57; Beckman Coulter, Brea, CA, USA; 1:100 dilution) in combination with Nef or Vpu rabbit antisera (1:1000 dilution). The percentage of infected cells (p24^+^) was determined by gating on the living cell population according to a viability dye staining (Aqua Vivid; Thermo Fisher Scientific). Alternatively, intracellular staining was assessed on 293T expressing Nef or Vpu proteins. Briefly, 2×10^6^ 293T cells were transfected with 7ug of Nef or Vpu expressor with the calcium-phosphate method. At 48 h post transfection, 293T cells were stained intracellularly with rabbit antisera raised against Nef or Vpu (1:1000). Samples were acquired on an LSRII cytometer (BD Biosciences), and data analysis was performed using FlowJo v10.5.3 (Tree Star, Ashland, OR, USA).

### FACS-based ADCC assay

Measurement of ADCC using the FACS-based assay was performed at 48h post-infection as previously described (43, 119). Briefly, HIV-1-infected primary CD4+ T cells were stained with AquaVivid viability dye and cell proliferation dye eFluor670 (Thermo Fisher Scientific) and used as target cells. Autologous PBMC effectors cells, stained with cell proliferation dye eFluor450 (Thermo Fisher Scientific), were added at an effector: target ratio of 10:1 in 96-well V-bottom plates (Corning, Corning, NY). A 1:1000 final dilution of plasma or 5μg/mL of A32 mAb was added to appropriate wells and cells were incubated for 5 min at room temperature. The plates were subsequently centrifuged for 1 min at 300 *x* g, and incubated at 37°C, 5% CO_2_ for 5h before being fixed in a 2% PBS-formaldehyde solution. Samples were acquired on an LSRII cytometer (BD Biosciences) and data analysis was performed using FlowJo v10.5.3 (Tree Star). The percentage of ADCC was calculated with the following formula: (% of p24+ cells in Targets plus Effectors) – (% of p24+ cells in Targets plus Effectors plus plasma) / (% of p24+ cells in Targets) by gating on infected lived target cells.

### Software Scripts and Visualization

Correlograms were generated using the corrplot package in program R v.4.1.012 and RStudio v.1.4.1106 (154, 155). Correlation networks were created using the ggraph and igraph packages in R in undirected mode, clustered based on the igraph layout “star”. Edges are weighted according to P-values (inversely). Edges are only shown if P < 0.05, and nodes without edges were removed. Nodes are sized according to the r-values of connecting edges. Multiple linear regression analyses were performed using the GraphPad Prism software (version 9.1.0). The coefficient of determination (R^2^) was used as a metric to measure the proportion of the variation observed with the dependant variable that can be explained by the variation in the independent variables. Since R^2^ values usually increases when more predictive variables are added to the model, we also measured the adjusted R^2^ (adj. R^2^) to account for this caveat.

### Statistical analysis

Statistics were analyzed using GraphPad Prism version 9.1.0 (GraphPad, San Diego, CA, USA). Every data set was tested for statistical normality and this information was used to apply the appropriate (parametric or nonparametric) statistical test. P values <0.05 were considered significant; significance values are indicated as * P<0.05, ** P<0.01, *** P<0.001, **** P<0.0001.

## Notes

### Competing Interest Statement

The authors have declared no competing interest.

